# A single N^1^- methyladenosine on the large ribosomal subunit rRNA impacts locally its structure and the translation of key metabolic enzymes

**DOI:** 10.1101/313874

**Authors:** Sunny Sharma, Johannes David Hartmann, Peter Watzinger, Arvid Klepper, Christian Peifer, Peter Kötter, Denis LJ Lafontaine, Karl-Dieter Entian

**Author notes:** Corresponding author: Correspondence should be addressed to Prof. Dr. Karl-Dieter Entian and Dr. Sunny Sharma. Joint first author (Sunny Sharma, Johannes David Hartmann).

## Abstract

The entire chemical modification repertoire of yeast ribosomal RNAs and the enzymes responsible for it have recently been identified. Nonetheless, in most cases the precise roles played by these chemical modifications in ribosome structure, function and regulation remain totally unclear. Previously, we demonstrated that yeast Rrp8 methylates m^1^A_645_ of 25S rRNA in yeast. Here, using mung bean nuclease protection assays in combination with quantitative RP-HPLC and primer extension, we report that 25S/28S rRNA of *S. pombe*, *C. albicans* and humans also contain a single m^1^A methylation in the helix 25.1. We characterized nucleomethylin (NML) as a human homolog of yeast Rrp8 and demonstrate that NML catalyzes the m^1^A_1322_ methylation of 28S rRNA in humans. Our *in vivo* structural probing of 25S rRNA, using both DMS and SHAPE, revealed that the loss of the Rrp8-catalyzed m^1^A modification alters the conformation of domain I of yeast 25S rRNA causing translation initiation defects detectable as halfmers formation, likely because of incompetent loading of 60S on the 43S-preinitiation complex. Quantitative proteomic analysis of the yeast *Δrrp8* mutant strain using 2D-DIGE, revealed that loss of m^1^A_645_ impacts production of specific set of proteins involved in carbohydrate metabolism, translation and ribosome synthesis. In mouse, NML has been characterized as a metabolic disease-associated gene linked to obesity. Our findings in yeast also point to a role of Rrp8 in primary metabolism. In conclusion, the m^1^A modification is crucial for maintaining an optimal 60S conformation, which in turn is important for regulating the production of key metabolic enzymes.

---

Gene expression is a multistep process in which the genetic information is used to produce a functional gene product – either RNA or protein. The production and regulation of steady state levels of these gene products is the fundamental condition for cellular and organismal life. In case of protein coding genes the expression amounts are determined by several regulatory loops that go beyond either transcription or translation, including programmed turnover of messenger RNAs and proteins ^1,2^. Surprisingly, gene expression for protein coding genes has always been correlated with transcription and it became a common practice to use mRNAs amount as proxies for the concentration of the corresponding protein ^1^. Nevertheless, recent parallel transcriptome and proteome analyses have highlighted several discrepancies with respect to this correlation, thus highlighting the need to understand so far unconsidered regulatory steps beyond transcription ^1, 3-5^.

Ribosomes are highly conserved ribonucleoprotein complexes that synthesize cellular proteins ^6^. Eukaryotic ribosomes consist of 4 ribosomal RNAs (rRNAs) and around 80 ribosomal proteins (r-proteins) ^7,8^. It is the rRNA which is responsible for the key activities of ribosomes namely decoding, peptidyl transfer and peptidyl hydrolysis ^9,10^. Until recently, ribosomes have always been seen as homogeneous, constitutive protein synthesizing machinery, lacking any major contribution in regulating gene expression ^11-13^ Usually, the efficacy of translation is suggested to be determined either by features intrinsic to the mRNAs or facilitated by protein or RNA adaptors (translational factors) ^1,2,14^. However, in contrast to this view, several recent elegant studies have highlighted indubitable roles of ribosomes in gene regulation ^15-18^. Emerging data have strengthened the perception that the ribosome population in cells is heterogeneous by virtue of the different components including ribosomal proteins, rRNAs and their chemical modifications. It becomes more and more evident that this ribosomal heterogeneity provides a further regulatory level of translational ^19^.

Synthesis of the two unequal subunits (40S and 60S) of the eukaryotic ribosome takes place in a highly regulated multi-step process comprising four rRNAs (25S/28S, 18S, 5.8S and 5S), 79 r-proteins in yeast (80 r-proteins in human), more than 200 non-ribosomal factors and 75 snoRNAs ^7^. The 18S and 25S rRNAs serve as the catalytic core of the ribosome, making it a “ribozyme” ^7,9^. They undergo various co- and post-transcriptional modifications during ribosome biogenesis. The most abundant chemical rRNA modifications are snoRNA guided methylation of the 2’-OH ribose moieties (Nm) and isomerization of uridines to pseudouridine (Ψ). These site specific modifications are guided by box C/D (Nm) and box H/ACA (Ψ) small nucleolar ribonucleoprotein particles (snoRNPs), respectively ^20,21^. Besides snoRNA-dependent rRNA modifications distinct rRNA base residues are modified by single enzymes ^21,22^. The 18S rRNAs of *Saccharomyces cerevisiae* contain seven base modifications, four base methylations (mN), two base acetylation and one amino-carboxypropylation ^23-27^. The complete set of base modified residues has been reported recently (2 m^1^A, 2 m^5^C and 2 m^3^U m^1^acp^3^Ѱ) ^28-31^. Most base modifications are either introduced at positions close to the peptidyl transferase center (PTC, m^1^A_645_, m^5^C_2870_ & m^3^U_2634_) or close to the subunit interface (m^1^A_2142_ & m^5^C_2278_) ^21^. Identification of the cellular machinery involved in chemical modifications is absolutely critical to decipher their role in the functioning of the ribosome and more importantly in cellular physiology.

Previously, we mapped the m^1^A_645_ modification in the helix 25.1 of the large subunit 25S rRNA in *Saccharomyces cerevisiae*, and identified the corresponding m^1^A-methyltransferase, Rrp8 ^28^. In the present study, we report that m^1^A in the helix 25.1 of the LSU rRNA is highly conserved in eukaryotes and demonstrate that in human cells it is catalyzed by Nucleomethylin (NML), a human homolog of yeast Rrp8. Furthermore, using RNA structure probing (DMS and SHAPE) and DIGE (Differential in gel electrophoresis), we demonstrated that m^1^A plays a critical role in maintaining proper 25S rRNA structure and that its loss affects profoundly the steady-state-levels of key metabolic enzymes.

## Results

### N^1^-methyladenosine in helix 25.1 of 25S/28S is a highly conserved base modification

The budding yeast, *S. cerevisiae* contains a single N^1^-methyl adenosine modified residue (m^1^A_645_) in helix 25.1 of the 25S rRNA ^28^. Alignment of 25S/28S rRNA sequences from *Candida albicans, Schizosaccharomyces pombe* and *Homo sapiens* revealed a highly-conserved sequence pattern around this modified m^1^A residue (Figure 1A). Using mung bean protection assays (Figure 1B) we isolated 25/28S rRNA fragments corresponding to the helix 25.1 of *C. albicans, S. pombe* and colon carcinoma (HCT116) human cell lines. RP-HPLC analyses of these fragments revealed that like *S. cerevisiae*, helix 25.1 of 25/28S rRNA in these organisms contain a single m^1^A residue, specifically: *C. albicans* (m^1^A_643_), *S. pombe* (m^1^A_670_) and *H. sapiens* (m^1^A_1322_) (Figure 1B).

**Figure 1.**
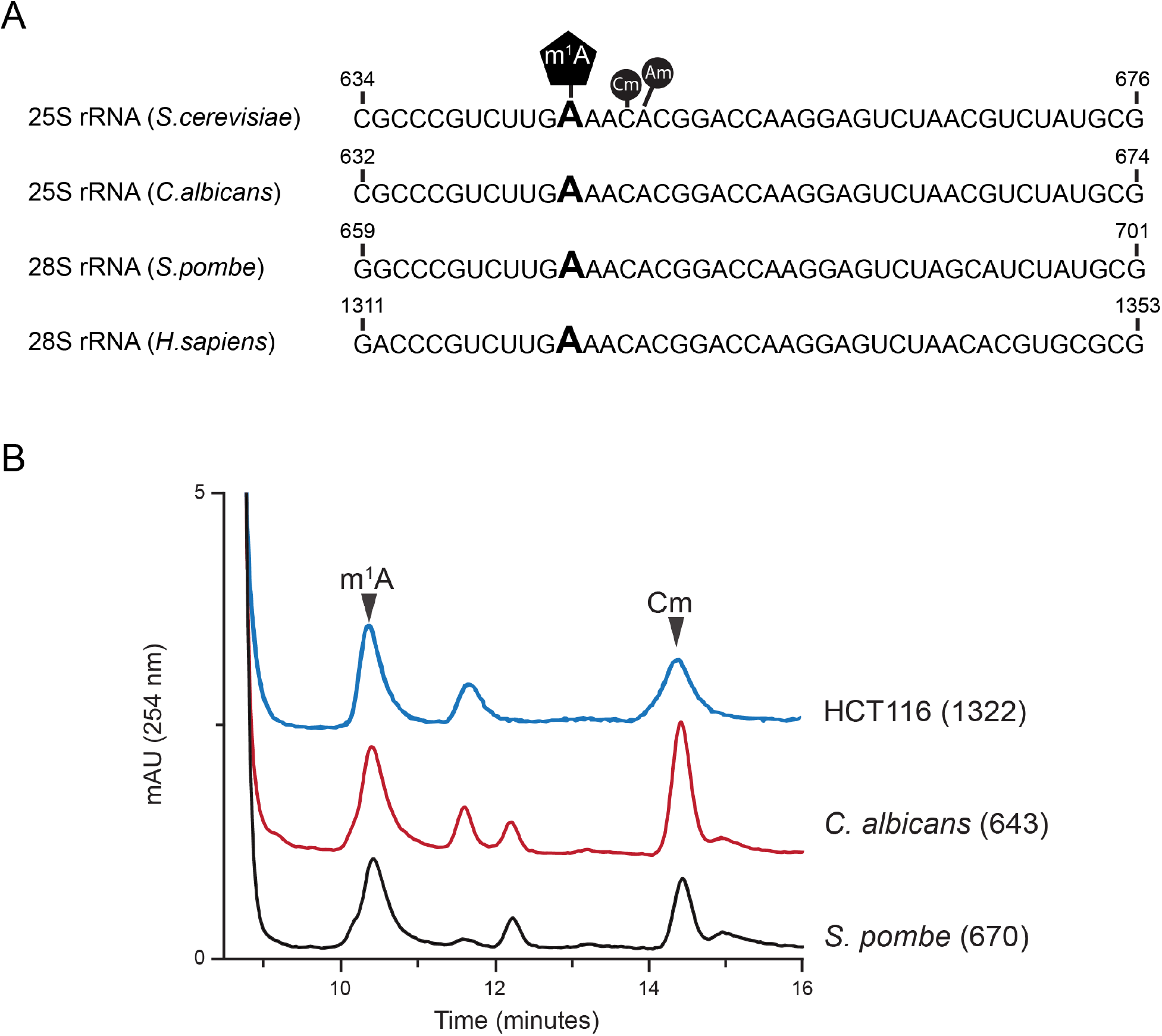
Conservation of the *N*^1^-methyladenosine in helix 25.1 of 25/28S rRNA. (A) rRNA sequence alignment of helix 25.1 from *S. cerevisiae*, *C. albicans, S. pombe* and *H. sapiens*. The m^1^A modified residue is highly conserved within helix 25.1 (B) Overlaid RP-HPLC chromatograms of nucleosides derived from fragments isolated using mung bean nuclease assay, containing m^1^A_643_ (Ca-Oligo643) isolated from wild-type *C. albicans* strain RM1000, m^1^A_670_ (Sp-Oligo643) isolated from wild-type *S. pombe* strain CBS356 and m^1^A_1322_ (Hs-Oligo1322) isolated from HCT116 cells. Cm (2’-O-ribose methylated cytidine) and Am (2’-O-ribose methylated adenosine) peaks are also labelled.

### Nucleomethylin (NML) catalyzes the methylation of m^1^A_1322_ of 28S rRNA in human cell lines

Previously, we identified Rrp8 as an *S-adenosyl L-methionine* (SAM)-dependent methyltransferase responsible for catalyzing N^1^-methylation of A_645_ of the 25S rRNA in *S. cerevisiae* ^28^. Bioinformatics analysis of the amino acid sequences of Rrp8 using The Basic Local Alignment Search Tool (BLAST) revealed that Rrp8 is not only conserved in several yeast species including *C. albicans* and the evolutionary distant *S. pombe*, but share a significant homology with the human Nucleomethylin (NML; Figure S1), especially in the C-terminal region. To validate that these Rrp8 homologs from *C. albicans* (*CaRRP8*) *S. pombe* (*SpRRP8*) and *H. sapiens* (*HsRRP8*/*NML*) function as m^1^A MTase, the respective genes were heterologously expressed in a *ScRRP8* deletion mutant (*Δrrp8*). Using mung bean protection assays and primer extension analyses, the methylation competences of all homologs were analyzed as explained in Materials and Methods. *CaRRP8* as well as *SpRRP8* showed 83.3 and 85.7 % methylation of A_645,_ respectively, whereas, even if over-expressed the human homolog, NML methylated A_645_ only to an extent of about 30 % compared with the *S. cerevisiae* wild-type (Figure 2A, 2B and 2C)

**Figure 2.**
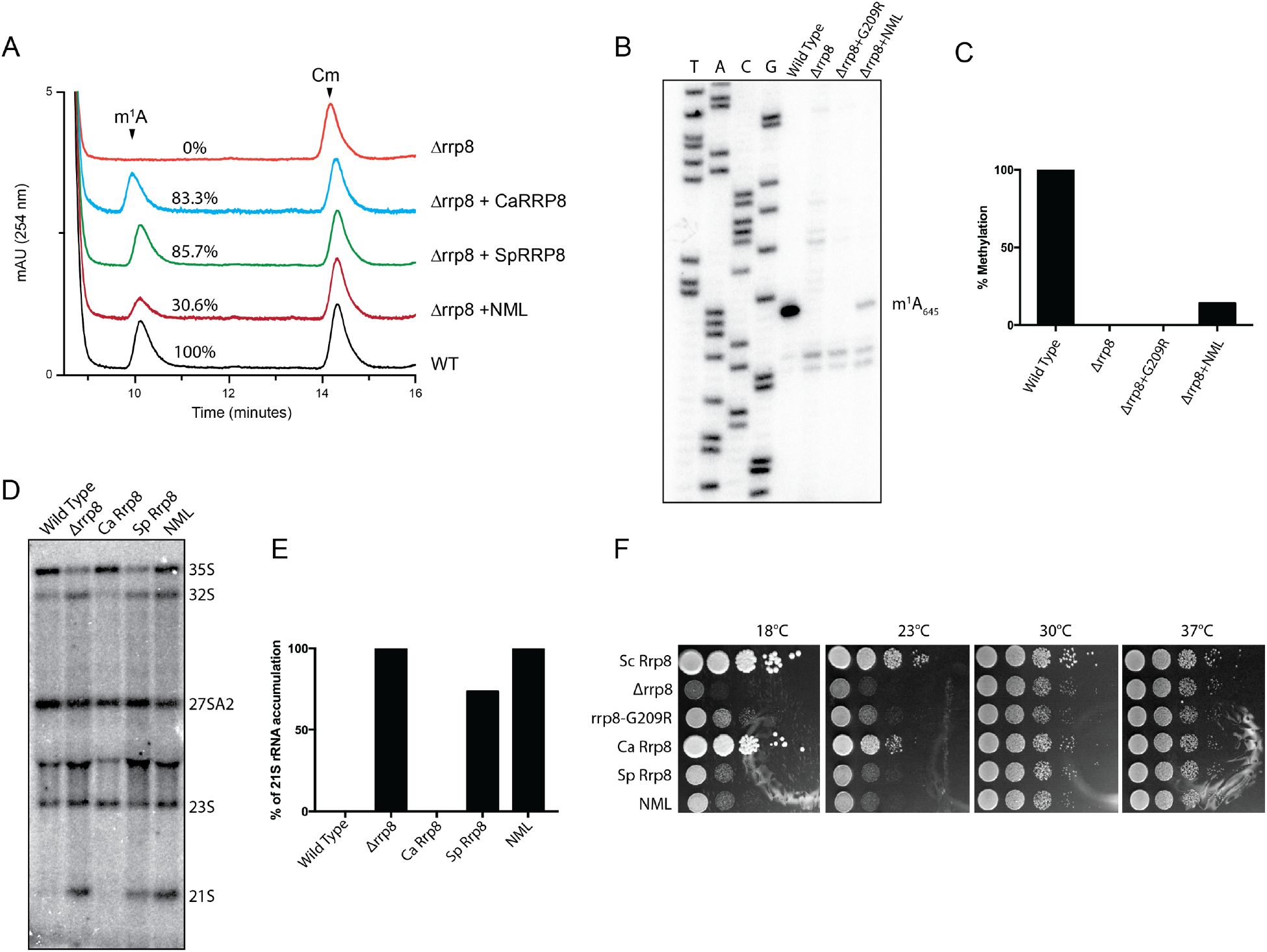
Heterologous expression of RRP8 homologs and biochemical and phenotypic analysis. (A) Overlaid RP-HPLC chromatograms of the nucleosides derived from fragments isolated using mung bean nuclease assay, containing m^1^A_645_ (Sc-Oligo645) isolated from wild-type (WT), deletion mutant (Δ*rrp8*) or deletion mutant strains expressing homologs of RRP8 (CaRRP8 for *C. albicans* Rrp8, SpRRP8 for *S. pombe* Rrp8 and NML for *H. sapiens* Rrp8/NML). (B) Primer extension analysis of Δ*rrp8*, the loss of methylation mutant rrp8^G209R^ and the heterologous expressed human homolog NML. The presence of m^1^A at position 645 in the 25S rRNA from the wild-type cells led to a strong stop. The expression of the human homolog NML in the deletion mutant of RRP8 of *S. cerevisiae* also led to a stop at this position to an extent of about 40% compared with *S. cerevisiae* wild-type. (C) Percent methylation was quantified from the intensity of primer extension stop (panel B) using ImageJ software (http://imagej.nih.gov/ij/). (D) Northern blot analysis of *RRP8* homologs expressed in *S. cerevisiae*. 5 μg of isolated total RNA was separated on an 1.2 % agarose-gel and subsequently blotted onto a Hybond H+ nylon membrane. The membrane was hybridized with radioactive (^32^P)-labeled probes against the internal transcribed spacer 1 (ITS1) (region between cleavage sites A_2_-A_3_). (E) Percent of 21S rRNA accumulation in panel D was quantified using ImageJ software. (F) Ten-fold dilutions of strains expressing RRP8, RRP8 homologs or catalytically-defective mutants. Cells spotted onto solid YEPD plates were incubated at four different temperatures illustrating the cold temperature sensitive phenotype of Δ*rrp8*.

Loss of Rrp8 leads to an accumulation of the aberrant 21S pre-rRNA due to defects in A_2_ cleavage and to cold sensitivity for growth ^32^. Interestingly, both of these phenotypes were fully complemented by *CaRRP8,* whereas the expression of *SpRRP8* could not restore 21S accumulation defects and cold sensitivity (Figure 2D, 2E and 2F). The human NML failed to complement the pre-rRNA processing phenotype and the growth cold sensitivity of *Δrrp8* (Figure 2D and 2F).

Having demonstrated that human NML is able to methylate yeast 25S rRNA *in vivo*, we next analyzed the impact of an NML knock down on m^1^A_1322_ methylation in HCT116 cells (Figure 3A and 3B). Therefore, these human cells were transfected for 72 hours with three different siRNAs (#166, #167 and #168), targeting distinct regions of the NML mRNA in three independent knock down experiments. The efficacies of NML depletions were analyzed both at the mRNA levels by RT-qPCR and at the protein levels by immunofluorescence, using commercially available antibody specific to NML protein (Figure 3B and 3C). All three siRNAs (#166, #167 and #168) led to a substantial depletion of the steady-state-levels of NML mRNA and consequently of protein (Figure 3C). Our immunofluorescence analysis also revealed that NML is primarily localized in the nucleolus and as observed in Figure 3C, depletion of NML leads to abnormal and altered nucleolar morphology.

**Figure 3.**
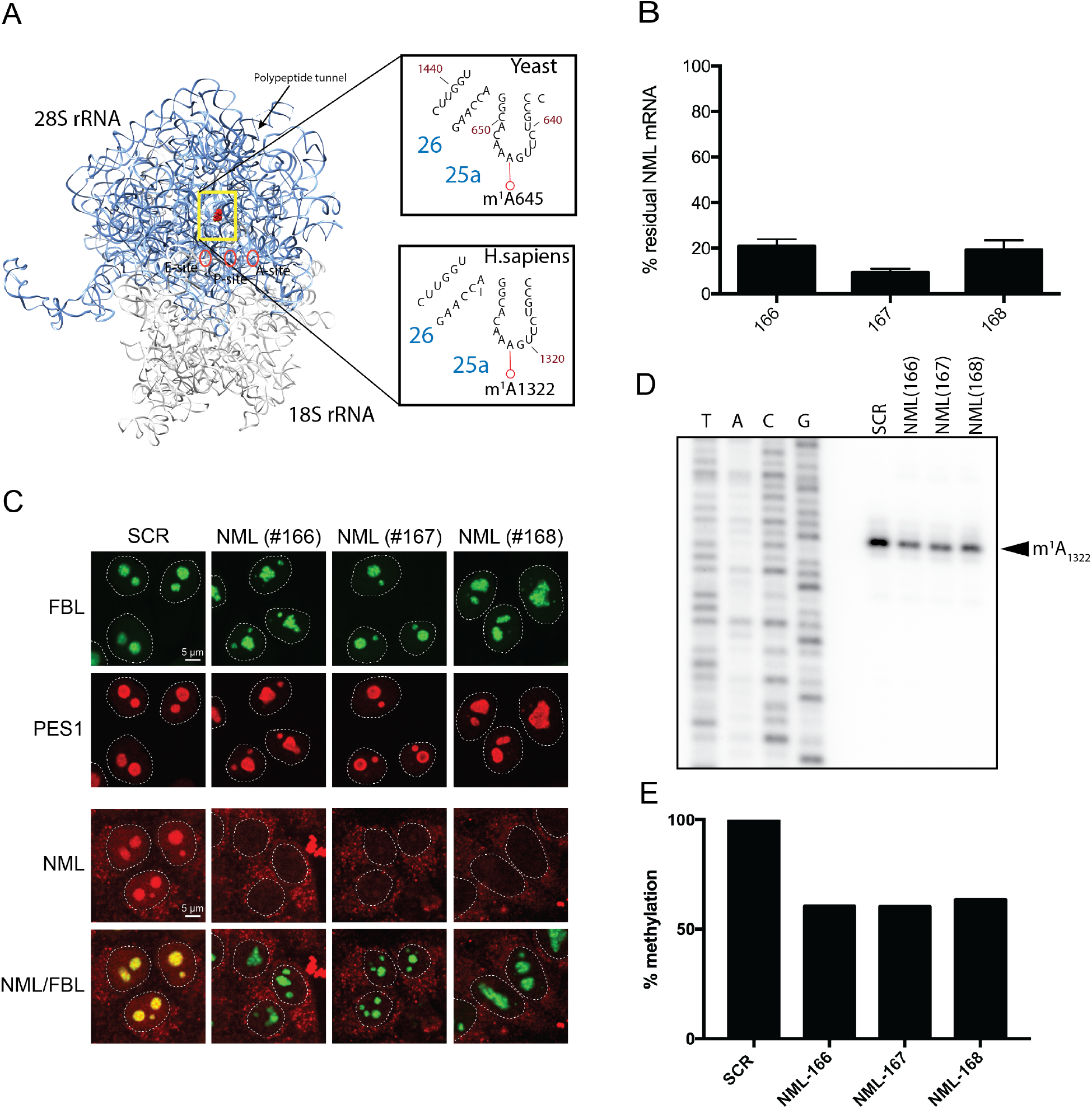
Validation of nucleomethylin (NML) as the human m^1^A large subunit rRNA methyltransferase. (A) 3-D structure model of the human rRNA. 4UG0 pdb file together with UCSF-Chimera were used to create the rRNA surface model. The N1-methylated residue A1322 in the 28S rRNA is highlighted as a red sphere. For comparison, the corresponding region of the yeast 25S rRNA is shown in an inset. (B) siRNA-mediated depletion of NML is highly efficient at mRNA level. Total RNA extracted from HCT116 cells transfected with a siRNA specific for NML (siRNA #166 to #168) and incubated for 72 h, was tested by RTqPCR. (C) NML is a nucleolar protein. Immunofluorescence with an antibody against NML was performed in HCT116 cells stably expressing a green fluorescent fibrillarin construct, which labels the dense fibrillar component of the nucleolus. The granular component protein PES1 was detected with a specific antibody (Bethyl Lab Inc). (D) Primer extension analysis of m^1^A methylation after siRNA-mediated depletion of NML in HCT116 cells. ^32^P-labeled primer complementary to nucleotides 1362 to 1381 of human 28S rRNA were used for methylation analysis of m^1^A_1322_. (E) The reduction of stop intensity at the m^1^A_1322_ in siRNA-mediated depleted cells was quantified using Image J software and estimated to be of ~60%.

Next, using primer extension analysis we demonstrated that knock-down of NML leads to around 40 % decrease in m^1^A_1322_ methylation (Figure 3D and 3E). These results established that m^1^A in the helix 25.1 of 25S/28S rRNA is a highly conserved chemical modification and is catalyzed by Rrp8 homolog, NML in human cells.

### Loss of m^1^A_645_ in yeast alters the rRNA topology by affecting eL32 interaction with 25S rRNA

Methylation of the N^1^ atom of adenosine leads to a net positive charge on the base and is expected to disrupt the canonical Watson-Crick base pairing ^33^. These chemical properties of m^1^A enhance non-canonical hydrogen bonding and electrostatic interactions, potentially promoting changes in RNA topology ^34,35^. To analyze any putative role of m^1^A_645_ on rRNA structure we performed *in vivo* dimethyl sulfate (DMS) and “selective 2’-hydroxyl acylation and primer extension” (SHAPE) RNA structure probing using our previously characterized catalytically-dead mutant of rrp8 ^G209R^ (28). DMS is among one of the most versatile chemical probe that can directly donate a methyl group to specific hydrogen bond accepting ring nitrogens on A, C, and G residues in RNA. The efficiencies of the methylation reaction depend on the chemical milieu of each base. Poor solvent accessibility and secondary structures protect from DMS methylation ^36,37^. SHAPE analysis using 2-methylnicotinic acid imidazolide (NAI), on the other hand, modifies all four nucleobases by acylation of the 2’-hydroxyl of flexible or accessible nucleotides mainly available within single-stranded RNA regions ^37,38^. Using DMS and SHAPE probing, we tested whether the loss of m^1^A_645_ leads to structural changes in and around the helix 25.1 region of the 25S rRNA, we primarily focused our analysis on 25S rRNA domain I. As a control for specificity, we also probed helix 72 of 25S rRNA with both reagents.

Indeed, both DMS and SHAPE rRNA structure probing revealed that upon loss of m^1^A_645_, regions proximal to helix 25.1 undergo a significant change in conformation, especially around expansion segments ES7b and ES7c (Figure 4 and 5A). As proof of specificity, no such structural alterations were observed in helix 72 (Figure S2). Mapping the residues for which sensitivity to DMS or NAI is altered upon loss of methylation on the 3-D structure of 60S revealed an interesting pattern of the structural changes in domain I of 25S rRNA. As illustrated in Figure 5B, N terminal residues of eL32 physically interact with the helix 25.1, particularly the D39 of eL32 likely establishes an electrostatic interaction with a positively charged m^1^A_645_. As described recently, eL32 forms a neuron like network with five different ribosomal proteins – uL22/L17, eL33/L33, eL6/L6, eL14/L14, uL30/L7 and uL4/L4 of the large subunit as shown in Figure 5A ^39^. It appears that the loss of m^1^A_645_ disrupts the interaction of helix 25.1 with eL32 that causes an allosteric effect on the whole domain by altering its conformation, mediated likely by a “synaptic network” of ribosomal proteins. This is well supported by an increase in reactivity of the region of the rRNA that directly interacts with these neighboring proteins (Figure 5A).

**Figure 4.**
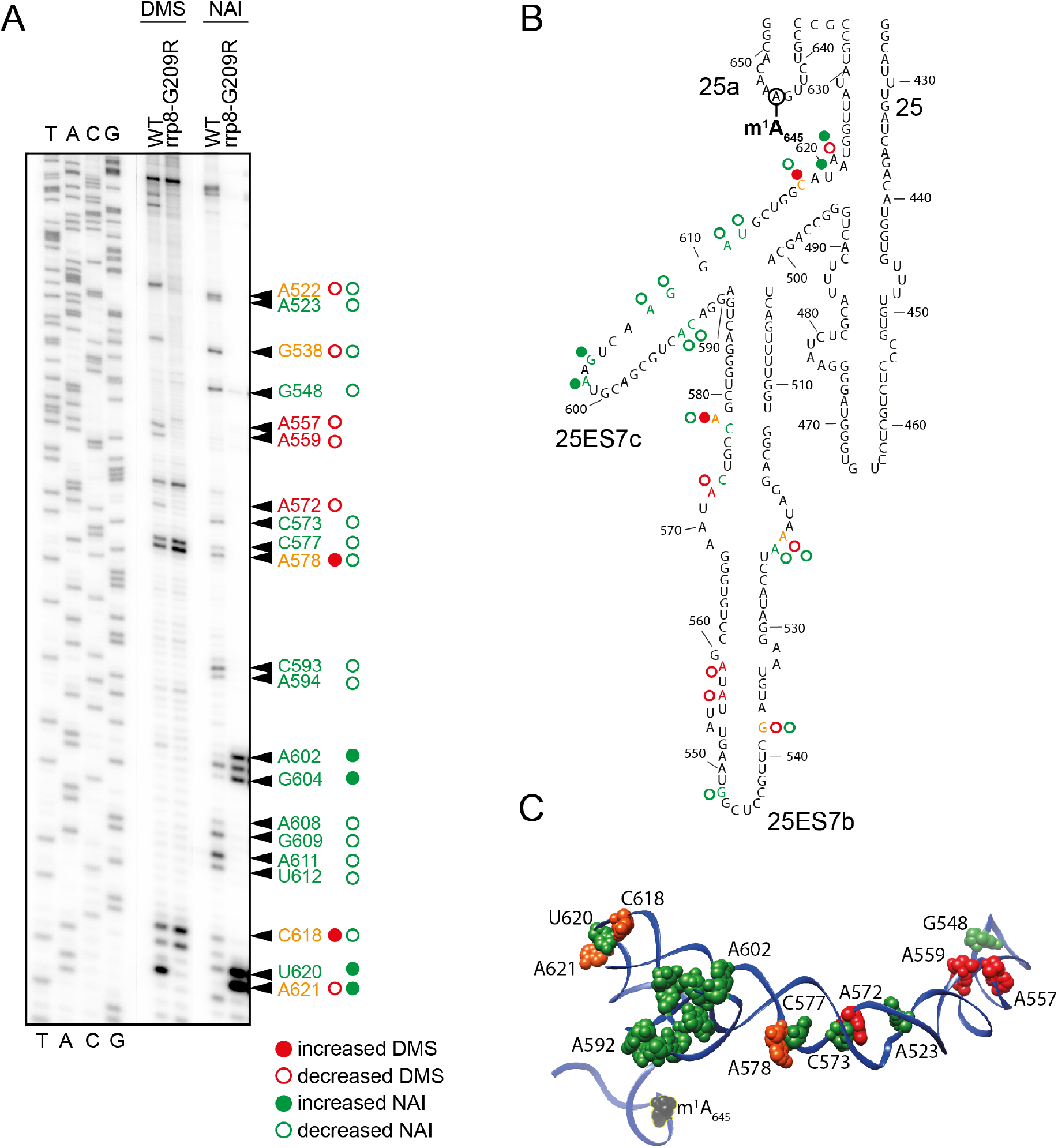
Loss of m^1^A_645_ leads to structural changes within the LSU. (A) Representative gel showing structure probing with DMS or SHAPE analysis with NAI in the rrp8^G209R^ loss of methylation mutant. ^32^P-labeled primer (helix25_StrPrb) complementary to nucleotides 2428 to 2448 of human 28S rRNA were used for the analysis. Bands corresponding to modified residues are marked in red for DMS, green for NAI and orange for both. (B) 2-D structure model with mapped modified residues as described in (A). (C) The residues sensitive to chemical modification are represented as spheres on a 3-D structure model of helix 25 (color scheme as in panel A). The model was made using Chimera and PDB file 4V88.

**Figure 5.**
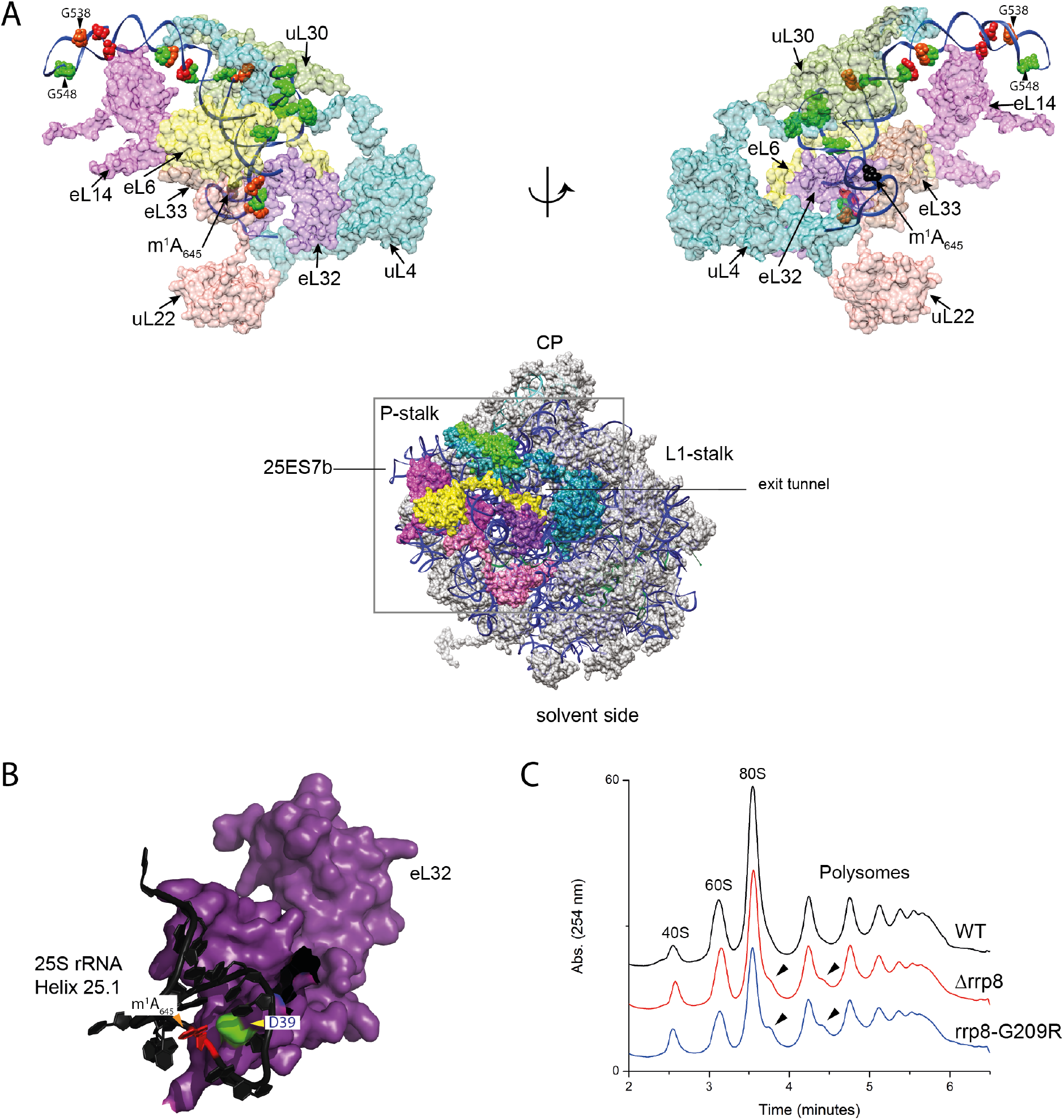
Loss of m^1^A_645_ leads to topological changes within the rRNA in the area of eL32 association (A) eL32 ‘neuron-like’ interaction network with five neighboring ribosomal proteins (uL22/L17, eL33/L33, eL6/L6, eL14/L14, uL30/L7 and uL4/L4) of the LSU around expansion segment ES7b and 7c. The models were made using Chimera and PDB files 4V88 (B) Possible electrostatic interaction of the D39 residue (green) of eL32 (colored violet) with the m^1^A_645_ modification (in red). 25S rRNA is colored black (C) Overlayed polysome profile analysis of *S. cerevisiae* wild-type (WT), Δr*rp8* and rrp8^G209R^ mutant strains. Polysome profiles of Δ*rrp8* and rrp8^G209R^ mutants were performed to detect the translational status in comparison with its isogenic wild-type. Half-mer formation in Δ*rrp8* and rrp8^G209R^ mutants are indicated by arrows.

Loss of m^1^A_645_ methylation causes formation of stalled preinitiation complex called halfmers (Figure 5C), which represent 43S initiation complex blocked on the initiation codon AUG, awaiting for the binding of 60S subunit to form 80S (monosome) ^28^. Halfmers formation may be due to in translation initiation inhibitions, or to defects in 60S assembly altering the stoichiometry of 40S-to-60S ratio ^40^. Considering that loss of m^1^A_645_ does not affect 60S synthesis and cause wide spread conformational changes that we observed in the present study using DMS and SHAPE probes, we speculate that 60S lacking m^1^A_645_ are less competent to bind to 40S subunits and therefore causes the formation of halfmers in the absence of m^1^A_645_.

### Quantitative protein expression analysis by 2D-DIGE

Having established that loss of m^1^A_645_ leads to local structural alterations in 25S rRNA domain I, we were interested to learn what the impact of such conformational alteration might have on translation. To address this, we performed whole proteome analysis by 2D-DIGE (2 Dimensional-Differential In Gel Electrophoresis) in cells lacking the 25S rRNA m^1^A_645_ modification and isogenic wild-type control cells.

Whole-cell extracts were prepared from three independent cultures of wild-type (CEN.PK2-1C) and *Δrrp8* mutant cells, pre-labeled with spectrally resolvable fluorescent dyes, either Cy3 (*Δrrp8* mutant) or Cy5 (wild-type control). To normalize for differences in protein abundance, a ‘mixed pool’ containing equal amounts of mutant and wild-type samples was labeled with the fluorescent dye Cy2. The pooled internal standard represents the average of all compared samples. Samples independently labelled with Cy2, Cy3, or Cy5, representing the proteins derived from the different conditions were mixed and resolved on the same 2-D gels. For each gel analyzed, we observed highly reproducible patterns of fluorescent protein spots (see Figure 6A for an example). We only considered for further analysis, the proteins which formed a clear spot after labeling with Cy3 and Cy5. Altogether, a total of ~1900 proteins were detected (~1,400 for pH 4- 7 and ~500 for pH 6-11) and quantitated. Proteins with statistically significant 1.5-fold differences in abundance between the wild-type and *Δrrp8* samples (95 % confidence interval as determined by Student *t*-test) were considered as significant. Comparing the wild-type and the *Δrrp8* mutant patterns revealed that the large majority of proteins were expressed to a similar extent. However, eighteen protein spots showed >1.5-fold differences. Of these, three proteins were upregulated in the *rrp8* mutant, whereas a decreased expression was found for fifteen proteins (Figure 6A, 6B and Table 1).

**Figure 6.**
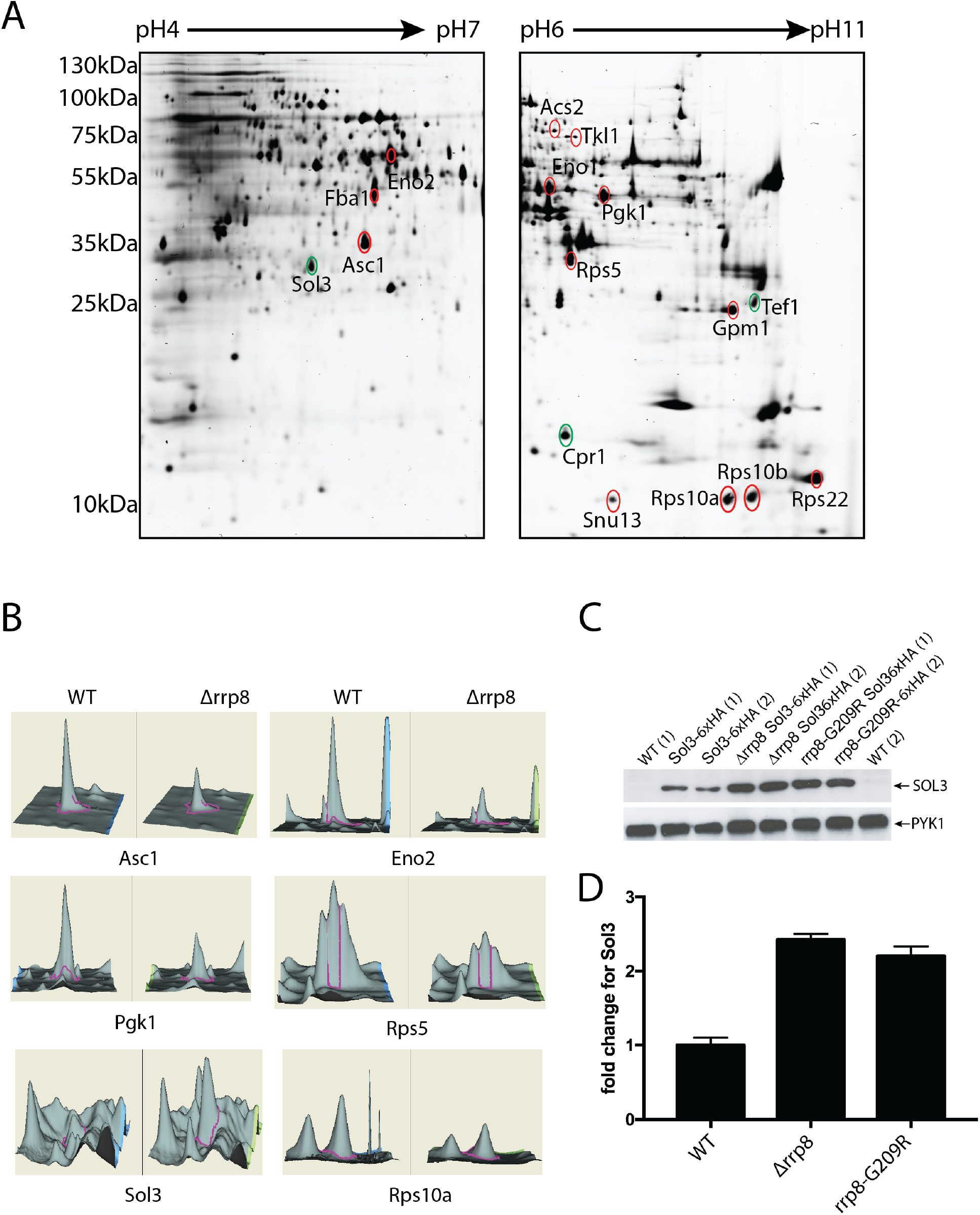
2D proteome analysis of rrp8 mutant. (A) Scanned images of a typical 2-D DIGE gel of 50 μg protein from the Cy3 labeled protein pool used for the rrp8 mutant strain (left gel: pH4-7, right gel: pH 6-11). Differentially expressed proteins are annotated and were identified using ESI-MS/MS. Differentially expressed protein spots are annotated in red (down-regulated) or green (up-regulated). (B) DeCyder output showing the representative differentially expressed protein spots of Asc1, Eno2, Pgk1, Rps5, Sol3 and Rps10A along with their 3-D fluorescence intensity profiles. (C and D) Sol3 up-regulation is related to loss of m^1^A_645_ methylation. Western blot analysis using anti-HA antibodies to analyze the expression of 6xHA tagged SOL3 in wild-type, Δ*rrp8* and rrp8^G209R^ mutant cells. As loading control, PYK1 was detected with a specific antibody. The represented fold changes in SOL3 expression were quantified using ImageJ software (http://imagej.nih.gov/ij/).

**Table 1.** Identification of differentially expressed proteins in a *RRP8* deletion strain compared to its isogenic wild-type control.

| **protein name** | **accession no.** | **p-value** | **no. of matched peptides** | **score** | **sequence coverage** | **fold change** |
| --- | --- | --- | --- | --- | --- | --- |
| **ENO2** | gi\|6321968 | 0.008 | 9 | 889 | 33 % | **-1.86** |
| **FBA1** | gi\|486081 | 0.014 | 4 | 515 | 25 % | **-1.86** |
| **RHR2** | gi\|1236750 | 0.0043 | 3 | 303 | 27 % | **-1.77** |
| **ASC1** | gi\|6323763 | 0.0033 | 1 | 55 | 6 % | **-1.87** |
| **SOL3** | gi\|338810340 | 1.1e<sup>-005</sup> | 10 | 91 | 40% | **+7.26** |
| **ACS2** | gi\|1360586 | 0.00051 | 2 | 160 | 8 % | **-1.6** |
| **TKL1** | gi\|1314142 | 0.034 | 5 | 439 | 10 % | **-1.63** |
| **ENO1** | gi\|1502357 | 0.0086 | 2 | 223 | 9 % | **-1.56** |
| **PGK1** | gi\|14588927 | 0.033 | 4 | 424 | 27 % | **-1.6** |
| **ADH1** | gi\|1419926 | 0.00022 | 6 | 667 | 26 % | **-1.88** |
| **RPS5** | gi\|1015849 | 0.018 | 2 | 268 | 19 % | **-1.54** |
| **TEF1** | gi\|32693293 | 0.011 | 7 | 144 | 18% | **+1.98** |
| **GPM1** | gi\|207343624 | 0.039 | 16 | 757 | 56% | **-1.57** |
| **RPS22A** | gi\|49258827 | 0.0084 | 8 | 290 | 64% | **-1.78** |
| **RPS10B** | gi\|6323886 | 0.016 | 4 | 412 | 57% | **-1.53** |
| **RPS10A** | gi\|536534 | 0.0022 | 3 | 227 | 40 % | **-1.57** |
| **SNU13** | gi\|6320809 | 0.005 | 5 | 294 | 56% | **-1.61** |
| **CPR1** | gi\|6320359 | 0.0068 | 7 | 302 | 50% | **+1.67** |

Surprisingly, after identification by mass spectrometry, the majority of the regulated proteins were found to correspond to enzymes involved in carbon metabolism and to proteins required for ribosome biogenesis and/or function. Additionally, acetyl-CoA-synthetase 2 (Acs2), which together with its isoenzyme Acs1 provides among others the nuclear source of acetyl for histone acetylation, was 1.6-fold decreased, whereas Cpr1, a cytoplasmic peptidyl-prolyl *cis-trans* isomerase increased approximately 1.7-fold (Table 1).

The largest gene ontology group that showed a significant difference in expression in the *Δrrp8* mutant, as compared to the wild-type, included nine proteins involved in carbon metabolism. Of these, six are glycolytic proteins, namely: fructose-1,6- bisphosphate aldolase, 3-phosphoglycerate kinase, phosphoglycerate mutase, enolase I and II and alcohol dehydrogenase I, two are pentose phosphate enzymes, and the last one is Gpp1, the glycerol-3-phosphate phosphatase, which generates glycerol needed to balance the redox potential and to adapt to anaerobic and osmotic stress. All glycolytic enzymes were decreased to a similar range comprised between 1.56 and 1.86-fold. It is important to note that although such variations may appear as marginal at first, one should bear in mind that because glycolytic proteins are so abundant in cells (corresponding to no less than ~5-10 % of total protein), even modest alterations will have major repercussions in terms of overall production of the protein of interest. The most remarkable alteration of expression we observed was an increase of 7-fold for the Sol3 protein (Figure 6, Table 1). Sol3 catalyzes the second step in the oxidative pentose phosphate pathway encoding 6-phosphogluconolactonase ^41,42^. To further analyze and validate the effect of loss of function of Rrp8 on Sol3 expression, we engineered strains that express a chromosomally-encoded HA-tagged Sol3 either in the rrp8 deletion (*Δrrp8*) or rrp8-catalytically deficient mutant (rrp8^G209R^). Protein expression analysis via Western blotting showed that Sol3 is not only upregulated in the *Δrrp8* mutant but that, importantly, it is more abundant in the rrp8^G209R^ mutant (Figure 6C). Thus, the catalytically deficient Rrp8 mutant (rrp8^G209R^) expresses ribosomes whose structure is locally perturbed in domain I of 25S rRNA, and cells harboring them are subjected to similar regulation of gene expression as those deleted for *RRP8*, at least as far as Sol3, our best candidate, is concerned. In both cases, Sol3 is upregulated at the protein level >2 fold, whereas mRNA levels remained unaffected (Figure 7A) suggesting that ribosomes lacking m^1^A_645_ translates Sol3 mRNAs more efficiently. Note that we believe the extent of fold change variation observed in the 2D-DIGE and Western blotting (7-fold and 2-fold increase for Sol3, respectively) simply reflects the difference in sensitivity of the assays used (the fluorescent signal from in vivo-labelled proteins is stronger than that resulting from Western blot detection by chemiluminescence). Finally, another enzyme of the pentose phosphate pathway with changed expression was Tkl1 that connects this pathway to glycolysis. As observed for all glycolytic enzymes, Tkl1 was also considerably decreased (Table 1).

**Figure 7.**
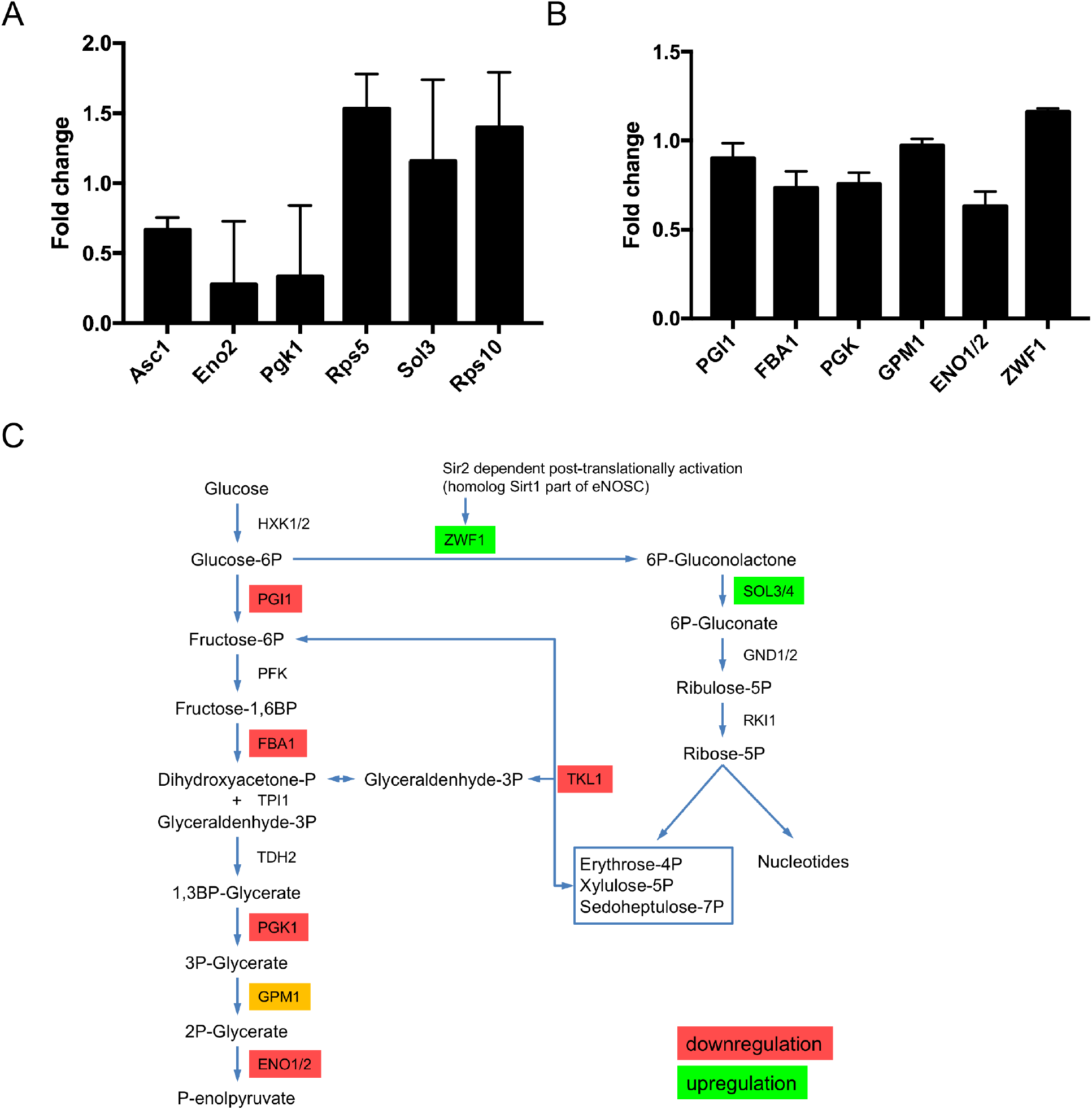
Quantitative PCR and biochemical activities of differentially regulated proteins in Δ*rrp8*. (A) RTqPCR analysis of mRNAs from proteins that were differentially regulated in the rrp8 mutant strain are represented as fold change in mRNA expression. Expression of Asc1, Rps5, Eno2 and Pgk1 is regulated at the transcriptional level, whereas mRNA levels of Sol3 and Rps10A are not different from their wild-type level of expression. Data analyzed with the software REST®.41(B) Regulation of specific activities of enzymes of the carbohydrate metabolism within Δ*rrp8*. Changes in the specific enzyme activities of phosphoglucose isomerase (PGI1), fructose-1,6- bisphosphate aldolase (FBA1), phosphoglycerate kinase (PGK1), phosphoglycerate mutase (GPM1), enolase (ENO1/ENO2) and glucose-6-phosphate dehydrogenase (ZWF1) are represented as fold change. Activities of enzymes involved in the glycolytic pathway are downregulated due to deletion of RRP8 as compared to wild-type. Only activity of ZFW1, first enzyme of the pentose-phosphate pathway, is slightly upregulated. (C) Schematic overview of regulations in the carbohydrate metabolism in cells with inactive RRP8.

To strengthen the 2-D DIGE observations and to better assess the physiological impact of the gene expression alterations on cell physiology, we measured the biochemical activities of 6 glycolytic enzymes (Figure 7B, and Materials and Methods section for details on the individual assays). Globally, we observed that the biochemical activity of the enzymes tested was altered to an extend compatible with the decreased amounts of proteins. Eno1 and Eno2 as well as Pgk1 showed reduced activities in agreement with the mRNA and protein reduction. Also, Fba1 showed decreased activity supporting the results of the 2D-DIGE experiments. Slightly decreased activities were monitored for Pgi1 and Gpm1 further sustaining a general downregulation in glycolysis (Figure 7B and 7C).

To evaluate a possible upregulation of the pentose phosphate pathway, the enzyme activity of Zwf1 (glucose-6-phosphate dehydrogenase) was measured. Glucose-6- phosphate dehydrogenase, which catalyzes the first step in this pathway ^43^, also showed a clear increase in enzyme activity (Figure 7B and 7C). A significant upregulation of Sol3 protein levels as depicted by our 2D-gel, emphasizes a possible induction of the pentose phosphate pathway, perhaps either as a consequence of downregulated glycolysis or facilitating the downregulation in glycolysis.

Apart from the proteins involved in carbohydrate metabolism, proteins involved in ribosome biogenesis and function also showed differential expression in the *Δrrp8* mutant. Identified proteins of this functional group are ribosomal proteins of the small subunit Rps5, Rps22, Rps10a and Rps10b and Asc1. These proteins displayed a 1.53- 1.87-fold downregulation. The decrease in core proteins of the 40S subunit and the downregulation of RNA binding protein that is part of the U3 snoRNP (Snu13) is in agreement with previous findings that showed Rrp8 is involved in pre-rRNA processing at site A_2_, resulting in the accumulation of aberrant rRNA precursors. The downregulation of ribosomal proteins might therefore be due to defects in ribosome assembly. Yeast Asc1 and mammalian RACK1 are functionally orthologous ribosomal proteins of the 40S subunit of which yeast Asc1 is reported to regulate expression of Eno2 which is also downregulated in the *Δrrp8* mutant ^44^. The *Δrrp8* deletion also caused an increase in protein abundance of the translational elongation factor EF-1 a (Tef1) as a possible compensation of a decreased ribosome amount.

### Analysis of steady state levels of mRNA of differential expressed proteins in a *Δrrp8* mutant

To further investigate the level of regulation (transcriptional or post-transcriptional) for the specifically identified proteins, we compared 2-D DIGE protein abundance data with mRNA levels by performing RT-qPCR experiments for several candidates. The protein abundance level from 2D-DIGE analysis and the mRNA levels established by RT-qPCR provided interesting insights (Figure 7A). Whereas the amount of the 40S subunit protein Rps5p was decreased at the protein level, the mRNA level was surprisingly upregulated, suggesting a post-translational regulation in the mutant. We additionally observed an upregulation of the Rps10A mRNA level. The transcriptional regulation of these ribosomal proteins suggests that the excess of mRNAs is either not translated or the proteins are post-translationally degraded. The mRNA levels of Asc1, Hri1, Acs2, Eno2 and Pgk1 correspond to their protein levels, suggesting that these proteins were likely regulated at the level of transcription.

## Discussion

Ribosomal RNAs are universally modified and rRNA modifications often cluster in functionally important areas of the ribosome ^21,45^. The complete chemical modification repertoire of yeast rRNAs and the cellular machinery responsible for it have recently been described ^21,27^ placing us in a position to address the putative roles of these chemical modifications in ribosome structure, function and regulation. Here, we have addressed the involvement in translation of a conserved m^1^A modification on the large ribosomal subunit and showed it is important to maintain locally the structure of the subunit with repercussions on the translation of specific transcripts encoding key metabolic enzymes.

There are only two known m^1^A modifications on yeast ribosomes both are located on the 60S subunit ^28,29^. The m^1^A modification we chose to study in this work is located on residue A645 of helix 25 on 25S rRNA, where it is deposited by the Rossmann-fold methyltransferase Rrp8. Strikingly, m^1^A_645_ is contacted by ribosomal protein eL32 (Figure 5A-B and ^21,28^). The functional implications of this proximity had not been addressed. Interestingly, *Rrp8* deletion in yeast cells leads to accumulation of unmethylated large ribosomal subunit and results in cryosensitive growth phenotype (impaired growth at the low temperatures of 18°C and 23°C). Furthermore, they are characterized by a mild pre-rRNA processing defect (visible by the accumulation of an aberrant 21S RNA reflecting altered kinetics of early nucleolar cleavages) without gross inhibition of mature subunit production (Figure 2).

We have shown that m^1^A modification in the helix 25.1 of 25S/28S is a highly-conserved chemical modification and is conserved throughout the eukaryotic kingdom being present in fission yeasts (*S. pombe*, Sp), in pathogenic yeasts (*C. albicans,* Ca), and in human cells (colon carcinoma) (Figure 1). We have validated the human nucleolar protein *nucleomethylin* (NML) as a structural and functional homolog of *RRP8* showing it is responsible for human 28S rRNA m^1^A methylation in the equivalent rRNA domain, domain 25a. We also report that NML is required to optimally maintain nucleolar integrity (Figure 3).

With respect to the impact of m^1^A_645_ modification on helix 25 and ribosome function, we made three significant observations in yeast cells: first, we showed that loss of m^1^A leads to local structural alterations in 25S rRNA at the sites of contacts of half a dozen of highly connected r-proteins, including eL32 (Figures 4-5). Atomic resolution structures of yeast ribosomes as shown that these r-proteins physically interact with each other at multiple instances forming an intricate ‘neuronal’ network ^39^. Second, we report that cells lacking the m^1^A modification accumulate halfmers. Since the formation of mature subunits is not grossly affected in these cells, we conclude that the accumulation of halfmers reflects translational initiation defects (at the step of 60S subunit joining). Third, our whole cell proteome analysis by 2-D DIGE, and the assessment of selected mRNA levels by RT-qPCR, have revealed that the expression of several key metabolic enzymes is substantially altered in cells lacking RRP8 (*Δrrp8* cells), as well as in cells lacking the m^1^A modification (rrp8^G209R^) (Figure 6). The most striking effect was seen for the 6-phosphogluconolactonase Sol3 (7-fold increase in expression) (Figure 7C).

In conclusion, we have demonstrated in budding yeast cells that removing a single base modification on the large ribosomal subunit is sufficient to specifically impact the translation of subsets of mRNA transcripts, in particular, those encoding key metabolic enzymes. We believe our observation provide a rationale for understanding why *nucleomethylin*, the mammalian homolog of RRP8, had previously been linked to nutrient availability signaling pathways and to obesity.

## Methods

### Bioinformatics analysis

All multi-alignments were performed with the open-source software T-coffee ^46^. EsPript was used to visualize alignments ^47^. Chimera (USF) was used to produce 3-D models of ribosomal subunits.

### Yeast strains and plasmids

All strains and plasmids used in the present study are listed in Supplementary Table S1 & S2. Polymerase chain reaction primers used for construction of the plasmids are listed in Supplementary Table S3. Yeast homologs of RRP8 have been amplified from genomic DNA of *C. albicans* and *S. pombe*. NML was amplified from human cDNA. Plasmids pPK468-CaRRP8, pPK468-SpRRP8 and pPK468-NML were constructed using the gap-repair strategy in budding yeast. Introduction of point mutants was carried out using polymerase chain reaction (PCR) site-directed mutagenesis with Single-Primer Reactions IN Parallel (SPRINP) with high fidelity Pfu-DNA polymerase (Promega) ^48^.

### Growth conditions and yeast media

Yeast strains were grown at 30°C or 16°C in YEPD (1 % w/v yeast extract, 2 % w/v peptone, 2 % w/v glucose) or in synthetic drop out medium lacking uracil (0.5 % w/v ammonium sulfate, 0.17 % w/v yeast nitrogen base, 2 % w/v glucose).

### Polysome profiles

Polysome profiles were performed as described before ^25^. The yeast strains were grown in 100 mL YEPD medium at 30°C and 16°C to an OD_600_ = 1.0. Cycloheximide was added to a concentration of 100 μg/ml and cells were further incubated for 30 min. The cell culture was then harvested by centrifugation at 4°C and further on washed two times with 10 mL buffer A (20 mM HEPES/KOH pH 7.5, 10 mM KCl, 1.5 mM MgCl_2_, 1 mM EGTA and 1 mM DTT). After resuspension in 0.5 mL buffer A and addition of an equal volume of glass beads, cells were disrupted by intensive shaking. Equivalent amounts of cleared cell lysate were layered onto a 10-50 % (w/v) sucrose gradient (made using Gradient Master 107 (Biocomp)) and centrifuged at 19,000 rpm for 17 h at 4°C in a SW40 rotor using a Beckman ultracentrifuge (L-70; Beckman). Absorbance profiles at 254 nm were then analyzed in ISCO UA-5 absorbance monitor.

### Yeast drop test assay

Yeast drop test assays were performed as previously described ^28^. Serial dilutions of yeast strains grown over night were first diluted to an OD_600_ of 1 and then serially diluted 1:10. 5 μl of each culture and dilution was spotted onto a SCD-uracil plate and incubated at 16°C, 23°C, 30°C and 37°C.

### RNA extraction

Yeast total RNA extraction using phenol/chloroform was exactly performed as previously described ^49^.

Human cells total RNA extraction using Tri-reagent solution (Life Technologies) was performed as previously described ^50^.

### Reverse Phase-High Pressure Liquid Chromatography (RP-HPLC) and mung bean nuclease protection assay

RP-HPLC analysis as well as the mung bean nuclease protection assay were performed as described previously ^51^. Synthetic deoxyoligonucleotides complementary to the specific sequence of C633 – G680 of yeast 25S rRNA were used for protection by hybridization to the rRNA. After digestion by mung bean nuclease (MBN Kit: M0250S, NEB) and 0.05 mg/ml RNase A (Sigma-Aldrich) purification of protected fragments from 8 M urea-PAGE (13 %) was carried out by passive elution with 0.3 M NaAc by rotation at 4°C overnight. Precipitation of eluted rRNA fragments was done using 100 % EtOH. Isolated fragments were digested with nuclease P1 and bacterial alkaline phosphatase (Sigma Aldrich) and subsequently the nucleosides were analyzed by RP-HPLC on a Supelcosil LC-18-S HPLC column (25 cm × 4.6 cm × 5 μm) equipped with a precolumn (4.6 cm × 20 mm) at 30°C on an Agilent 1200 HPLC system.

### Human cell culture and siRNA-mediated depletion

Human cells (HCT116 p53 +/+) were grown at 37 °C under 5 % CO_2_. The cell lines used in this work were obtained directly from ATCC and passaged in the laboratory for fewer than 6 months after receipt. All ATCC cell lines were diagnosed by short tandem repeat (STR) profiling. HCT116+/+ cells were reverse transfected as follows: 1.5 μl of 20 μM siRNA (Life Technologies) and 4 μl Lipofectamine RNAiMAX (Life Technologies) were mixed with 500 μl Opti-MEM (Life Technologies) in each well of a 6-well plate. After a 20-min incubation at room temperature, 1.5 × 10^5^ cells (for 72 hours), were re-suspended in 2.5 ml antibiotic-free medium and were seeded into each well. The siRNAs (Life Technologies-Silencer Select) used are listed in Supplementary Table S3.

### Western-blotting

Western blot analysis was carried out using 25 μg of total protein extracts from HA-epitope tagged yeast strains. Protein samples were separated with 12 % sodium dodecyl sulphate (SDS-) PAGE and subsequent blotting on a PVDF membrane (Millipore). The membrane was blocked with 5 % nonfat dry milk. For the detection of tagged proteins anti-HA monoclonal antibody (Roche; 1:1000 dilution) were used following incubation with anti-mouse-IgG-conjugated horseradish peroxidase (BioRad; 1:10000 dilution). Pyk1 was detected using polyclonal anti-Pyk1(1:10000) followed by anti-rabbit-IgG-conjugated horseradish peroxidase (BioRad: 1:10000 dilution).

### Quantitative reverse transcription (RTqPCR)

Prior to the reverse transcription total RNA (~1 μg) was digested with DNase I (Sigma-Aldrich) according to the manufacturerŕs protocol. The DNase I digested RNA was then reverse transcribed using random hexamers and SuperScript II reverse transcriptase (Invitrogen) according to the manufactureŕs protocol. Real-time quantitative polymerase chain reactions were performed in a Rotor-Gene 3000 Real-Time Cycler (QIAGEN/Corbett) with 0.5 μl of the product from the reverse transcriptase reaction using gene specific primers (0.5 μl of 20 μM stock solution) and a Maxima^®^ SYBR Green/ROX qPCR Master Mix (Fermentas) in a final volume of 25 μl. Primer sequences are listed in Supplementary Table S3. The PCR program used was 96°C for 10 min, 50 cycles of 96°C for 30 s; 56°C for 30 s; 72°C for 30 s, and a final stage of 72°C for 5 min. Relative gene expression was determined by the Pfaffl-method to calculate the average PCR efficiency for each primer set, multiple serial dilution curves were performed. For determination of the CT values, the Rotor-Gene software version 6.1 (Corbett Research) was used. The obtained CT values and PCR efficiencies were used to calculate the relative gene expression normalized to control Act1 values using the Relative Expression Software Tool (REST^®^) ^52^.

### RNA electrophoresis

Analysis of high-and low-molecular-weight species of yeast rRNA species as well as analysis of high-molecular-weight-species of human was performed as described previously ^50^.

### Northern blotting

Northern blotting was performed as previously described for yeast rRNA and human rRNA ^25^.

### Primer extension

Primer extension analysis were performed exactly as described previously ^27^. The primers used for performing the analyses are listed in Supplementary Table S3.

### Immunofluorescence

Zeiss Axio Observer.Z1 microscope with a motorized stage, driven by MetaMorph (MDS Analytical Technologies, Canada) was used for all imaging. Images were captured in widefield mode with a 20 objective (Plan NeoFluar, Zeiss), a LED illumination (CoolLed pE-2) and a CoolSnap HQ2 camera. High-resolution images were captured in confocal mode using a Yokogawa spin-disk head and the HQ2 camera with a laser from Roper (405 nm 100 mW Vortran, 491 nm 50 mW Cobolt Calypso and 561 nm 50 mW Cobolt Jive) and a 40 objective (Plan NeoFluar, Zeiss). After 72 hours of siRNA mediated knock down of Rrp8, cells were first fixed in 2% of formaldehyde, washed in PBS and then blocked in PBS supplemented with 5 % of BSA and 0.3 % Triton X-100 for 1 hour at 25°C. Rabbit anti-Rrp8 Antibody AbVantage Pack (Bethyl Lab Inc.) was diluted in PBS supplemented with 1 % BSA, 0.3 % Triton X-100 and incubated with the cells overnight at 4°C. Cells were washed in PBS and incubated with a secondary Alexa Fluor 594 anti-rat antibody (1:1,000; Invitrogen) in PBS, 1 % BSA, 0.3 % Triton X-100 for 1 h. Cells were finally washed in PBS, treated with DAPI and imaged with the Zeiss microscope as described above.

### *In vivo* DMS and SHAPE structure probing

100 mL YPD media was inoculated with an overnight culture to a starting OD_600_ of 0.2 and grown at 30°C to an OD_600_ of 0.8 to 1.5. In the hood, two 15 mL aliquots of yeast culture in 50 mL polypropylene tubes were treated with 300 μL of 95% ethanol with 1:4, v/v DMS (Sigma) or without as a negative control and mixed vigorously for 15 seconds. Both aliquots were then incubated with shaking for 2 minutes at 30°C. The reaction was stopped by placing the tube on ice and adding 5 mL of 0.6 M β-mercaptoethanol and 5 mL of isoamyl alcohol. After addition of the stop solutions the tubes were vortexed for 15 seconds and then centrifuged at 3000×*g* at 4°C for 5 minutes. The cell pellets were washed with another 5 mL of 0.6 M β-mercaptoethanol.

Total RNA was isolated from the DMS treated cell pellet using GTC (4 M guanidium isothiocyanate, 50 mM Tris-HCl pH 8.0, 10 mM EDTA pH 8.0, 2% w/v sodium lauroyl sarcosinate and 150 mM β-mercaptoethanol) as described above. The sites of A and C methylation in by DMS were mapped by primer extension using primers binding proximal to the helix 25.1 (helix25.1_StrPrb and Str_Prb_h25b) and helix 72 (helix72_StrPrb). Primer extension was carried out exactly as described previously^27^. RNA SHAPE analysis were performed exactly as described ^37^ using the same primers as used for DMS probing.

### Proteome Analysis

#### Sample preparation and labelling

Yeast cells were harvested from a 100 ml logarithmic culture (OD_600_ = 2; grown at 30°C) and subsequently washed with water and then PGSK solution (50 mM NH_2_PO_4_, 4 mM Na_2_HPO_4_, 50 mM NaCl, 5 mM KCl, 60 mM glucose and 30 mM Tris-HCl pH 8.8). The cell pellet was re-suspended in 500 μl of PGSK-Buffer and an equal amount of glass beads were added. Cells were disrupted at 4°C for 5 minutes with 30 s incubation on ice after every minute. The tubes were centrifuged for 10 min at 3,000 g, followed by 15,000 g for 10 minutes and the supernatant was collected in a fresh cup. The protein concentration in the supernatant was determined using the Bradford Reagent (SERVA Electrophoresis GmbH) and the protocol provided by the manufacturer.

An equal amount of proteins (50 μg) from the wild-type and mutant were labelled with Cy3 and Cy5 respectively according to the manufactureŕs protocol (GE Healthcare). A mixed-sample internal control generated by pooling an equal amount of wild-type and *Δrrp8* mutant proteins were labeled with Cy2. The Cy2 standard was used to normalize protein abundances across different gels and to control gel-to-gel variation.

### Two-dimensional Difference Gel Electrophoresis (2-D DIGE)

The samples (labeled proteins) from the wild-type, mutant and mixed internal standard were dissolved in 250 μl of Rehydration-Buffer (7 M urea, 2 M thio-urea, 4 % CHAPS with 0.5 % DTT and 1.2 % de-streaking solution) and 13 cm IPG strips with pH range 4- 7 and 6-11 were rehydrated over night with the Rehydration-Buffer carrying the labelled proteome of the wild-type and the *Δrrp8* mutant along with the internal standard. For the first dimension, the Ettan IPGphor-3 (GE Healthcare) was used to carry out isoelectric focusing (IEF). IEF was performed using the protocol mentioned in the GE handbook for respective pH range. Before subjecting the IPG strips to second dimension, an equilibration with Equilibration-Buffer (6 M urea, 20 mM Tris-HCl pH 8.8, 2 % SDS, 20 % glycerol) containing 1 % DTT (freshly added) followed by Equilibration-Buffer containing 4 % idoacetamide (added fresh) for 20 min each were carried out. After equilibration, IPG strips were blotted between two moist filter papers and then placed horizontally on to a polyacrylamide gel (12 %) which was then sealed by 2 % agarose containing bromophenol blue to perform the second dimension. The gels were run at 4°C with constant current of 16 mA overnight using Biorad Protean II xi.

#### Image acquisition

The gels were scanned using the Typhoon 9400 Variable Mode Imager (GE Healthcare) with mutually exclusive emission and excitation wavelengths for each CyDye (GE Healthcare), generating overlaid multiple-channel images for each gel. The Cy2 images were scanned using a 468 nm laser and an emission filter of 520 nm BP (band pass) 40. Cy3 images were scanned using a 532 nm laser and an emission filter of 580 nm BP30. Cy5 images were scanned using a 633 nm laser and a 670 nm BP30 emission filter. The narrow BP emission filters reduce the cross-talk between fluorescent channels. All gels were scanned at 100 μm resolution. Images were initially analyzed and cropped using ImageQuant 5.2.

#### Image analysis

2-D DIGE analysis was performed with DeCyder version 6.5 software (GE Healthcare). All gel image pairs were initially processed by the DeCyder DIA module (differential-in-gel analysis) which uses a triple-co-detection algorithm to detect and generate the same protein spot-feature boundary for individual Cy2, Cy3, and Cy5 signal to calculate the Cy3/Cy2 and Cy5/Cy2 ratios. The images analyzed and processed in the DIA module were then later combined using the DeCyder BVA module (biological variation module) to compare the ratios among the four DIGE gels each for pH range 4-7 and pH range 6- 11. A Student t test analysis was applied for triplicate samples from all strains, despite separation on different DIGE gels. This provided the findings a greater statistical validity. In our analysis of *in vivo* protein levels, we considered 2-D gel features to be a protein only if the spot was confirmed with both Cy-labels and were present on every gel. Only proteins with 1.5-fold differences in abundance were considered significant. Differences in protein levels between wild-type and *Δrrp8* mutant samples were considered statistically significant only if the difference fell within the 95 % confidence interval as determined by the Student *t* test.

#### Protein identification

The selected 2-D gel spots were picked from the gel and mass spectrometry analyses were carried out at Proteomics Service of Applied Biomics, Inc. (Haywarth, CA, USA). Picked spots were washed twice with 25 mM ammonium bicarbonate and 50 % acetonitrile, once with water and once with 100 % acetonitrile to remove the staining dye and other inhibitory chemicals. Gel spots were then dried to absorb a maximum volume of digestion buffer. Dried 2-D spots were rehydrated in digestion buffer (25 mM ammonium bicarbonate, 2 % acetonitrile, containing 0.5 % sequencing grade modified trypsin (Promega, Madison, MI)). Proteins were digested in gel at 37°C. Digested peptides were extracted from the gel with TFA extraction buffer (0.1 % trifluoroacetic acid). The digested tryptic peptides were then desalted using C-18 Zip-tips (Millipore). The desalted peptides were mixed with CHCA matrix (alpha-cyano-4-hydroxycinnamic acid) and spotted into wells of a MALDI plates. Mass spectra of peptides in each sample were obtained using Applied Biosystems 4700 Proteomics Analyzer (Applied Biosystems, Foster City, CA). Proteins were identified based on peptide fingerprint mass mapping using MS spectra and peptide fragmentation mapping using MS/MS spectra. Combined MS and MS/MS spectra were submitted for database search using GPS Explorer software equipped with the MASCOT search engine to identify proteins from primary sequence databases. Only those 2-D gels spots were accepted as a positive identification which showed the highest proteins scoring hit with protein score confidence interval over 95 % from the database search.

### Enzyme Activity Assays

#### Preparation of cell extract

Cells harvested from 100 ml YEPD culture grown to an OD_600_ of 0.7 to 1 were re-suspended in 0.5 ml 100 mM potassium phosphate buffer (pH 6.5) or 100 mM sodium phosphate buffer (pH 6.6) and broken by heavy mixing with 500 μl glass beads. Glass beads and cell debris were separated by centrifuging for 10 min at 2600 g. The extract was kept on ice and directly analyzed.

#### Determination of protein concentration

The protein concentration was determined using the micro biuret method as described before ^53^. Therefore 100 μl cell extract, 500 μl micro biuret reagent (8 M NaOH, 0.21 % CuSO_4_) and 1 ml potassium phosphate buffer (pH6.5) or sodium phosphate buffer (pH6.6) were mixed and the absorption at 290 nm was measured using a DU 650 spectrophotometer (Beckman Coulter). Bovine serum albumin was used as standard.

#### Phosphoglucose isomerase (Pgi1)

The phosphoglucose isomerase activity was determined as described before ^43^ following the reduction of NAD^+^ in a coupled assay with glucose-6-phosphate dehydrogenase (G6P-DH) from *Leuconostoc mesenteroides*. The assay was conducted at 25°C in the following mixture: 50 mM TEA (pH 7.4); 10 mM MgCl_2_; 1 mM fructose-6-phosphate; 1 mM NAD^+^ and 2 U/ml G6P-DH. The use of G6P-DH from *Leuconostoc mesenteroides* thereby allowed for the use of NAD^+^ instead of NADP^+^. The assay was started by addition of fructose-6-phosphate to the assay mixture.

#### Fructose-1,6-bisphosphate aldolase (Fba1)

The fructose-1,6-bisphosphate aldolase activity was determined as described before ^43^ following the oxidation of NADH in a coupled assay with triose phosphate isomerase (TIM) from *S. cerevisiae* and glycerol-3- phosphate dehydrogenase (GDH) from rabbit muscle. The assay was conducted at 25°C in the following mixture: 50 mM TEA (pH 7.4); 10 mM MgCl_2_; 100 mM KAc; 1 mM fructose-1,6-bisphosphate; 0.1 mM NADH; 1 U/ml GDH and 10 U/ml TIM. The assay was started by addition of fructose-1,6-bisphosphate to the assay mixture.

#### Phosphoglycerate kinase (Pgk1)

The phosphoglycerate kinase activity was determined as described before ^43^ following the oxidation of NADH in a coupled assay with glycerinealdehyde-3-phosphate dehydrogenase (GAP-DH) from rabbit muscle. The assay was conducted at 25°C in the following mixture: 50 mM TEA (pH 7.4); 10 mM MgCl_2_; 1 mM 3-phosphoglycerate; 0.1 mM NADH; 1 mM ATP; 5 mM cysteine and 2 U/ml GAP-DH. It was started by addition of 3-phosphoglycerate to the assay mixture.

#### Phosphoglycerate mutase (Gpm1)

The phosphoglycerate mutase activity was also determined as described before ^43^ following the oxidation of NADH in a coupled assay with Enolase from *S. cerevisiae*, pyruvate kinase (PK) from rabbit muscle and lactate dehydrogenase (LDH) from hog muscle. The assay was conducted at 25°C in the following mixture: 50 mM TEA (pH 7.4); 10 mM MgCl_2_; 1 mM 3-phosphoglycerate; 0.7 mM NADH; 1 mM ADP; 1 mM 2,3-diphosphoglycerate; 1 U/ml Enolase; 2 U/ml PK and 2 U/ml LDH. It was started by addition of cell extract to the assay mixture.

#### Enolase (Eno1/Eno2)

The enolase activity was also determined as described before ^43^ following the oxidation of NADH in a coupled assay with pyruvate kinase (PK) from rabbit muscle and lactate dehydrogenase (LDH) from hog muscle. The assay was conducted at 25°C in the following mixture: 50 mM TEA (pH 7.4); 10 mM MgCl_2_; 1 mM 2-phosphoglycerate; 0.1 mM NADH; 1 mM ADP; 1 U/ml PK and 1 U/ml LDH. It was started by addition of 2-phosphoglycerate to the assay mixture.

#### Glucose-6-phosphate dehydrogenase (Zwf1)

The glucose-6-phosphate dehydrogenase activity was determined following the reduction of NADP^+^ in a direct assay. The assay was conducted at 25°C in the following mixture: 50 mM TEA (pH 7.4); 10 mM MgCl_2_; 1 mM glucose-6-phosphate and 0.3 mM NADP^+^. The assay was started by addition of glucose-6-phosphate to the assay mixture.

#### Enzyme unit definition

One unit was defined as the amount of protein necessary to convert 1 μmol of substrate per minute. All enzyme measurements were performed using a Specord S 600 spectrophotometer (Analytik Jena) following the absorption of NADH at 340 nm.

## References

1. Vogel, C. & Marcotte, E. M. Insights into the regulation of protein abundance from proteomic and transcriptomic analyses. Nat Rev Genet 5, 1512 (2012).

2. Gebauer, F. & Hentze, M. W. Molecular mechanisms of translational control. Nat. Rev. Mol. Cell Biol. 5, 827–835 (2004).

3. Tian, Q. et al. Integrated Genomic and Proteomic Analyses of Gene Expression in Mammalian Cells. Mol. Cell Proteomics 3, 960–969 (2004).

4. Vogel, C. et al. Sequence signatures and mRNA concentration can explain two-thirds of protein abundance variation in a human cell line. Molecular Systems Biology 6, 404 (2010).

5. Schwanhäusser, B. et al. Global quantification of mammalian gene expression control. Nature 473, 337–342 (2011).

6. Ramakrishnan, V. Ribosome structure and the mechanism of translation. Cell 108, 557–572 (2002).

7. Woolford, J. L. & Baserga, S. J. Ribosome biogenesis in the yeast Saccharomyces cerevisiae. Genetics 195, 643–681 (2013).

8. Henras, A. K., Plisson-Chastang, C., O’Donohue, M.-F., Chakraborty, A. & Gleizes, P.-E. An overview of pre-ribosomal RNA processing in eukaryotes. WIREs RNA 6, 225–242 (2015).

9. Nissen, P., Hansen, J., Ban, N., Moore, P. B. & Steitz, T. A. The structural basis of ribosome activity in peptide bond synthesis. Science 289, 920–930 (2000).

10. Polacek, N. & Mankin, A. S. The Ribosomal Peptidyl Transferase Center: Structure, Function, Evolution, Inhibition. Crit. Rev. Biochem. Mol. Biol. 40, 285– 311 (2005).

11. Deusser, E. & Wittmann, H.-G. Biological Sciences: Ribosomal Proteins: Variation of the Protein Composition in Escherichia coli Ribosomes as Function of Growth Rate. Nature 238, 269–270 (1972).

12. Byrgazov, K., Vesper, O. & Moll, I. Ribosome heterogeneity: another level of complexity in bacterial translation regulation. Curr Opin Microbiol 16, 133–139 (2013).

13. Xue, S. & Barna, M. Specialized ribosomes: a new frontier in gene regulation and organismal biology. Nat. Rev. Mol. Cell Biol. 13, 355–369 (2012).

14. Larsson, O. & Nadon, R. Gene expression - time to change point of view? Biotechnol. Genet. Eng. Rev. 25, 77–92 (2008).

15. Vesper, O. et al. Selective Translation of Leaderless mRNAs by Specialized Ribosomes Generated by MazF in Escherichia coli. Cell 147, 147–157 (2011).

16. Kondrashov, N. et al. Ribosome-Mediated Specificity in Hox mRNA Translation and Vertebrate Tissue Patterning. Cell 145, 383–397 (2011).

17. Schosserer, M. et al. Methylation of ribosomal RNA by NSUN5 is a conserved mechanism modulating organismal lifespan. Nat Commun 6, 6158 (2015).

18. Shi, Z. et al. Heterogeneous Ribosomes Preferentially Translate Distinct Subpools of mRNAs Genome-wide. Mol. Cell 1–21 (2017). doi:10.1016/j.molcel.2017.05.021

19. Simsek, D. & Barna, M. An emerging role for the ribosome as a nexus for post-translational modifications. Curr Opin Cell Biol 45, 92–101 (2017).

20. Decatur, W. A. & Fournier, M. J. rRNA modifications and ribosome function. Trends Biochem Sci 27, 344–351 (2002).

21. Sharma, S. & Lafontaine, D. L. J. ‘View From A Bridge’: A New Perspective on Eukaryotic rRNA Base Modification. Trends Biochem Sci 40, 560–575 (2015).

22. Sloan, K. E. et al. Tuning the ribosome: the influence of rRNA modification on eukaryotic ribosome biogenesis and function. RNA Biol 0–0 (2016). doi:10.1080/15476286.2016.1259781

23. Lafontaine, D., Delcour, J., Glasser, A. L., Desgrès, J. & Vandenhaute, J. The DIM1 gene responsible for the conserved m6(2)Am6(2)A dimethylation in the 3’- terminal loop of 18 S rRNA is essential in yeast. J Mol Biol 241, 492–497 (1994).

24. White, J. et al. Bud23 methylates G1575 of 18S rRNA and is required for efficient nuclear export of pre-40S subunits. Mol. Cell. Biol. 28, 3151–3161 (2008).

25. Sharma, S. et al. Yeast Kre33 and human NAT10 are conserved 18S rRNA cytosine acetyltransferases that modify tRNAs assisted by the adaptor Tan1/THUMPD1. NAR 43, 2242–2258 (2015).

26. Meyer, B. et al. Ribosome biogenesis factor Tsr3 is the aminocarboxypropyl transferase responsible for 18S rRNA hypermodification in yeast and humans. NAR 44, 4304–4316 (2016).

27. Sharma, S. et al. Specialized box C/D snoRNPs act as antisense guides to target RNA base acetylation. PLoS Genet 13, e1006804 (2017).

28. Peifer, C. et al. Yeast Rrp8p, a novel methyltransferase responsible for m1A 645 base modification of 25S rRNA. NAR 41, 1151–1163 (2013).

29. Sharma, S., Watzinger, P., Kötter, P. & Entian, K.-D. Identification of a novel methyltransferase, Bmt2, responsible for the N-1-methyl-adenosine base modification of 25S rRNA in Saccharomyces cerevisiae. NAR 41, 5428–5443 (2013).

30. Sharma, S., Yang, J., Watzinger, P., Kötter, P. & Entian, K.-D. Yeast Nop2 and Rcm1 methylate C2870 and C2278 of the 25S rRNA, respectively. NAR 41, 9062–9076 (2013).

31. Sharma, S. et al. Identification of novel methyltransferases, Bmt5 and Bmt6, responsible for the m3U methylations of 25S rRNA in Saccharomyces cerevisiae. NAR 42, 3246–3260 (2014).

32. Bousquet-Antonelli, C., Vanrobays, E., Gélugne, J. P., Caizergues-Ferrer, M. & Henry, Y. Rrp8p is a yeast nucleolar protein functionally linked to Gar1p and involved in pre-rRNA cleavage at site A2. RNA 6, 826–843 (2000).

33. Agris, P. F., Sierzputowska-Gracz, H. & Smith, C. Transfer RNA contains sites of localized positive charge: carbon NMR studies of [13C]methyl-enriched Escherichia coli and yeast tRNAPhe. Biochemistry 25, 5126–5131 (1986).

34. Macon, J. B. & Wolfenden, R. 1-Methyladenosine. Dimroth rearrangement and reversible reduction. Biochemistry 7, 3453–3458 (1968).

35. Mikhailov, S. N. et al. Chemical incorporation of 1-methyladenosine into oligonucleotides. NAR 30, 1124–1131 (2002).

36. Peattie, D. A. & Gilbert, W. Chemical probes for higher-order structure in RNA. Proc. Natl. Acad. Sci. U.S.A. 77, 4679–4682 (1980).

37. Spitale, R. C. et al. RNA SHAPE analysis in living cells. Nat Chem Biol 9, 18–20 (2012).

38. Tijerina, P., Mohr, S. & Russell, R. DMS footprinting of structured RNAs and RNA–protein complexes. Nat Protoc 2, 2608–2623 (2007).

39. Poirot, O. & Timsit, Y. Neuron-Like Networks Between Ribosomal Proteins Within the Ribosome. Sci. Rep. 6, 26485 (2016).

40. Helser, T. L., Baan, R. A. & Dahlberg, A. E. Characterization of a 40S ribosomal subunit complex in polyribosomes of Saccharomyces cerevisiae treated with cycloheximide. 1, 51–57 (1981).

41. Stanford, D. R. Division of Labor Among the Yeast Sol Proteins Implicated in tRNA Nuclear Export and Carbohydrate Metabolism. Genetics 168, 117–127 (2004).

42. Duffieux, F. Molecular characterisation of the first two enzymes of the pentose-phosphate pathway of trypanosoma brucei. J Biol Chem (2000). doi:10.1074/jbc.M004266200

43. Maitra, P. K. & Lobo, Z. A kinetic study of glycolytic enzyme synthesis in yeast. J. Biol. Chem. 246, 475–488 (1971).

44. Zeller, C. E., Parnell, S. C. & Dohlman, H. G. The RACK1 Ortholog Asc1 Functions as a G-protein beta Subunit Coupled to Glucose Responsiveness in Yeast. J Biol Chem 282, 25168–25176 (2007).

45. Lafontaine, D. L. J. Noncoding RNAs in eukaryotic ribosome biogenesis and function. Nature Structural & Molecular Biology 22, 11–19 (2015).

46. Di Tommaso, P. et al. T-Coffee: a web server for the multiple sequence alignment of protein and RNA sequences using structural information and homology extension. NAR 39, W13–7 (2011).

47. Gouet, P. ESPript/ENDscript: extracting and rendering sequence and 3D information from atomic structures of proteins. NAR 31, 3320–3323 (2003).

48. Edelheit, O., Hanukoglu, A. & Hanukoglu, I. Simple and efficient site-directed mutagenesis using two single-primer reactions in parallel to generate mutants for protein structure-function studies. BMC Biotechnol. 9, 61 (2009).

49. Yang, J., Sharma, S., Kötter, P. & Entian, K.-D. Identification of a new ribose methylation in the 18S rRNA of S. cerevisiae. NAR 43, 2342–2352 (2015).

50. Sharma, S., Marchand, V., Motorin, Y. & Lafontaine, D. L. J. Identification of sites of 2’-O-methylation vulnerability in human ribosomal RNAs by systematic mapping. Sci. Rep. 7, 11490 (2017).

51. Yang, J. et al. Mapping of Complete Set of Ribose and Base Modifications of Yeast rRNA by RP-HPLC and Mung Bean Nuclease Assay. PLoS ONE 11, e0168873 (2016).

52. Pfaffl, M. W., Horgan, G. W. & Dempfle, L. Relative expression software tool (REST) for group-wise comparison and statistical analysis of relative expression results in real-time PCR. NAR 30, e36 (2002).

53. Itzhaki, R. F. & Gill, D. M. A Micro-Biuret method for estimating proteins.Analytical Biochemistry 9, 401–410 (1964).

